# A high-precision hybrid algorithm for predicting eukaryotic protein subcellular localization

**DOI:** 10.1101/620179

**Authors:** Dahan Zhang, Haiyun Huang, Xiaogang Bai, Xiaodong Fang, Yi Zhang

**Author notes:** To whom correspondence should be addressed. Tel: []; Fax: []; Email: [ ], Correspondence may also be addressed to Xiaodong Fang. Tel: []; Fax: []; Email: [ ].

## Abstract

**Motivation:** Subcellular location plays an essential role in protein synthesis, transport, and secretion, thus it is an important step in understanding the mechanisms of trait-related proteins. Generally, homology methods provide reliable homology-based results with small E-values. We must resort to pattern recognition algorithms (SVM, Fisher discriminant, KNN, random forest, etc.) for proteins that do not share significant homologous domains with known proteins. However, satisfying results are seldom obtained.

**Results:** Here, a novel hybrid method “Basic Local Alignment Search Tool+Smith-Waterman+Needleman-Wunsch” or BLAST+SWNW, has been obtained by integrating a loosened E-value Basic Local Alignment Search Tool (BLAST) with the Smith-Waterman (SW) and Needleman-Wunsch (NW) algorithms, and this method has been introduced to predict protein subcellular localization in eukaryotes. When tested on Dataset I and Dataset II, BLAST+SWNW showed an average accuracy of 97.18% and 99.60%, respectively, surpassing the performance of other algorithms in predicting eukaryotic protein subcellular localization.

**Availability and Implementation:** BLAST+SWNW is an open source collaborative initiative available in the GitHub repository (https://github.com/ZHANGDAHAN/BLAST-SWNW-for-SLP or http://202.206.64.158:80/link/72016CAC26E4298B3B7E0EAF42288935)

**Supplementary Information:** Supplementary data are available at PLOS Computational Biology online.

## 1. INTRODUCTION

Following the completion of human, Arabidopsis thaliana [1], and rice genome sequencing projects in the early 21st century [2], additional investigations have been carried out in genome-related studies. Proteins transported to specific subcellular locations are guided by typical signals in protein sequences and secondary structures. Proteins sharing common primary and secondary structures are generally located in the same cellular regions [3]. Moreover, these features are typically used for subcellular localization prediction (SLP) [4]. The determination of subcellular localization (SL) is an arduous but indispensable task and is expected to lead to a thorough understanding of several areas, including signalling pathway, molecular mechanism, and individual development. However, even for well-studied organisms such as yeast [5,6], the efficiencies of numerous experimental protein-SL annotation methods, including bimolecular fluorescence complementation, Green fluorescent protein, and immunohistochemistry, are constrained by difficulties in antibody production, fusion protein preparation and active protein maintenance.

A series of machine learning methods and homology-based methods have been proposed to fill the gap in sequence-based SLP [7]. The most commonly used features in SLP include amino acid composition, including 20 amino acid frequency [8], dipeptide frequency [9], n-peptide frequency [10], pseudo-amino acid frequency [11], hydrophobic stretch, and indivisible N-terminal signal sequence [12]; functional domain composition [13] based on the function database SBASE-A and gene annotations (CELLO and YLOC) [14–16]. Other methods attempt to integrate and utilize multiple characteristics, including amino acid frequency and order information [17], functional domains, and pseudo-amino acid composition [18], higher-order dipeptide composition, N-terminal and C-terminal split amino acid composition and hybrid information [19], CTD (composition, translation, and distribution) of physiochemical descriptors and sequence similarity [20], signal peptides and targets [21], amino acid composition, N-terminal, and C-terminal features [22], sequence-based or text-based features [23].

Based on this information, SVM (20,21), N-to-1 neural networks [24], and Basic Local Alignment Search Tool (BLAST)-SVM union algorithms [25] have been widely utilized in SLP so far. Recently, a novel software package, LocTree3 [26], which integrates SVM with a homology algorithm, has been proposed to perform SLP on proteins from all domains of life, including Archaea, Bacteria, and Eukaryota. LocTree3 has been used in a number of recently published papers [27–29]. However, a recent study suggests that the performance of specifically developed machine learning tools shows no significant improvement over that of homology-search-based methods [30], which is consistent with what we demonstrate in this paper.

Here, a novel algorithm BLAST+SWNW is born by integrating BLAST with Smith-Waterman (SW) and Needleman-Wunsch (NW). In the first step of BLAST+SWNW, similar sequences to each test protein sequence are identified by BLAST (2.2.31) with an E-value=30 in the train dataset. Then, SW and NW are used to select the most similar sequence to the test protein sequence. Finally, the SL of the protein test sequence is predicted to be that of the most similar sequence judged by SW and NW. BLAST+SWNW integrates the high speed of BLAST with the high accuracy of SW and NW and offers a significant advantage over former SLP methods.

## 2. MATERIALS AND METHODS

### 2.1 Materials

As shown in **Fig 1**, two protein datasets (Dataset I and Dataset II) are constructed and used to test the performance of BLAST+SWNW and other algorithms.

**Fig 1.**
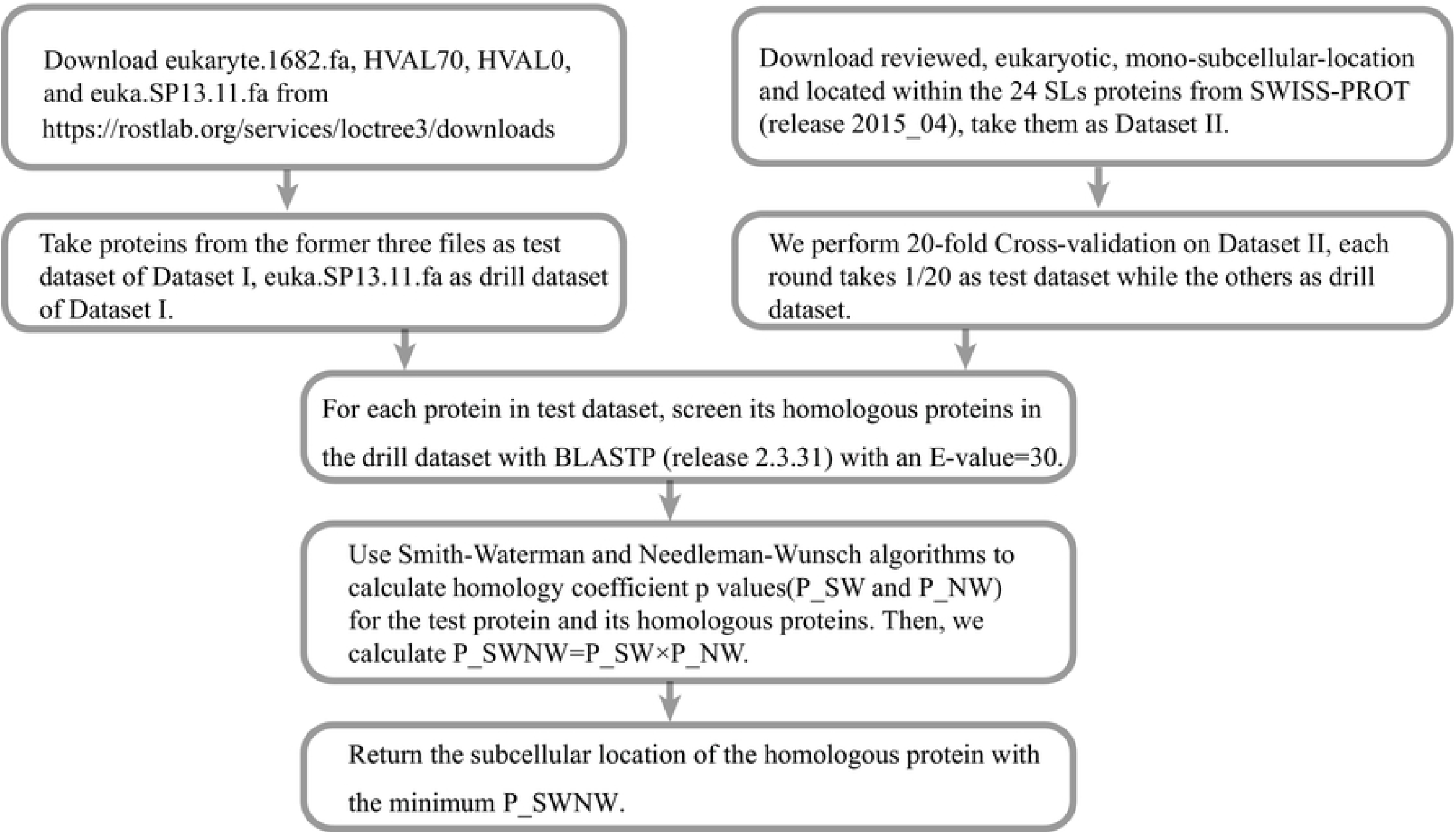
Flow chart of dataset construction and the BLAST+SWNW process.

All proteins in Dataset I are from LocTree3 Server [26] (https://rostlab.org/services/loctree3/downloads). Its 18-class-subcellular-location-coverage drill dataset includes 35,738 eukaryotic proteins from SWISS-PROT release 2013_11 (S1 Table), and its test dataset is composed of 3 independent test datasets T1, T2, and T3 that include 1682, 1053 and 273 eukaryotic proteins, respectively.

Dataset II proteins are from SWISS-PROT release 2015_04. First, unreviewed, multi-localization, missing item, or non-eukaryotic proteins were comprehensively removed from release 2015_04. Then, 24 SLs (**Fig 2** and **S2 Table**) were proposed as classifications for 110,168 reviewed mono-localization proteins. All data can be accessed at https://github.com/ZHANGDAHAN/subcellular-protein-prediction-data

**Fig 2.**
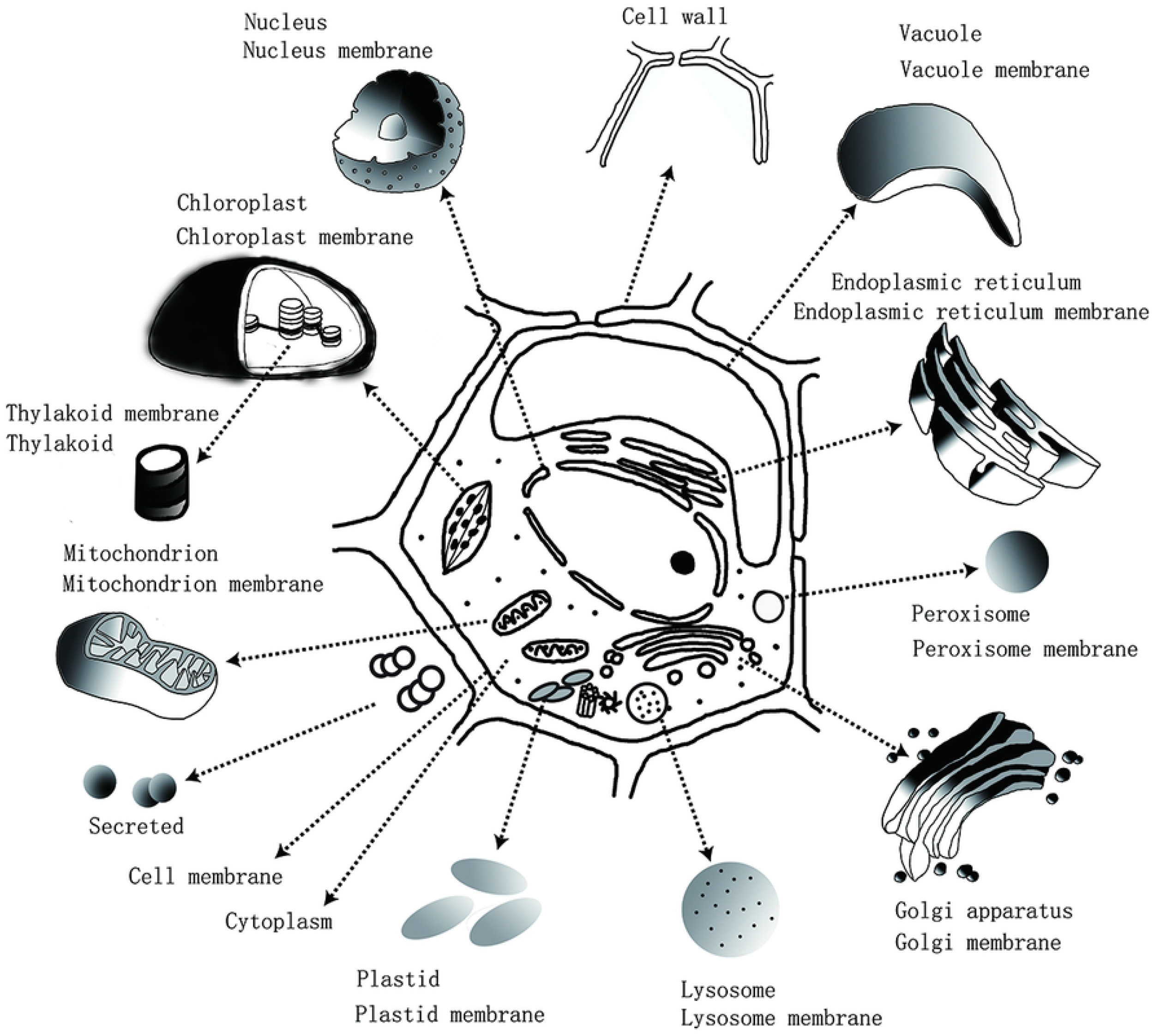
24 subcellular locations of Dataset II. This figure demonstrates 24 subcellular locations and their positions in an eukaryotic cell.

The 24 SLs in Dataset II consist of the 18 SLs proposed in Dataset I and another six novel SLs: “Cell wall”, “ Thylakoid “, “ Thylakoid membrane”, “ Lysosome”, “ Lysosome membrane”, and “Plastid membrane”. Proteins located in each SL generally play fundamental roles in cell biology. In detail, “Thylakoid” is an essential place for light-dependent reactions, consisting of the thylakoid membrane and thylakoid lumen [31]. “Lysosome” is a membrane-bound cellular organelle existing in most animal cells that harbours over 50 different enzymes responsible for plasma membrane repair, cell signalling, and energy metabolism [32]. “Plastid” has several differentiated forms and is subdivided into two SLs: “plastid” and “plastid membrane”. Proteins in the cell wall are key to structural support and morphogenesis, and their roles are nonnegligible [33].

### 2.2 Methods

Protein sequence homology comparison algorithms are effective methods for exploring proteins’ functions and predicting their SLs. In this paper, BLAST [34], SW [35] and NW [36] compose BLAST+SWNW.

#### 2.2.1 BLAST

BLAST is one of the best bioinformatics research tools, and its derivatives, such as BLASTP, PSI-BLAST, and PHI-BLAST, have been among the most widely used tools in alignment searches. BLASTP is a heuristic-based comparison tool [34] that uses seeded alignment heuristics and roughly compares entry sequences with a database to find similar sequences from it within a short time-frame. Therefore, we used BLASTP to screen similar sequences in the first step of BLAST+SWNW.

#### 2.2.2 Smith-Waterman and Needleman-Wunsch alignment algorithms

The SW algorithm is a dynamic programming algorithm [35]. SW compares segments of all possible lengths and optimize their similarity measures; hence, it is a precise algorithm to calculate the local alignment of two sequences. However, it is time consuming and space consuming as a query protein has to be compared with all sequences in a given protein database. The NW algorithm is a robust alignment program for optimal global alignment [36], but its penalty system differs from that of SW. Owing to their outperformance over others, these two algorithms have been integrated in BLAST+SWNW.

#### 2.2.3 The detailed process of BLAST+SWNW

We take Dataset I as an example to illustrate BLAST+SWNW (**Fig 1**). First, for a sequence (named “queryseq”) from the test dataset of Dataset I, we scan its similar protein sequences (named “databaseseqs”) from the drill dataset of Dataset I with BLASTP with a loosened E-value of 30. In this paper, 30 is the default Evalue of BLAST+SWNW. Second, we delete all non-standard amino acid characters from “queryseq” and “databaseseqs” to meet the input requirements of SW and NW. Third, p values between the “queryseq” and its similar “databaseseq” are calculated by integrating the output of SW and NW. In detail, for a “queryseq” and “databaseseq” pair, we set the parameter λ to 0.16931, k to 0.20441, and define m and n as the length of “queryseq” and “databaseseq”, respectively. Then, the similarity scores measured by SW (sw_score) and by NW (nw_score) are calculated using MATLAB functions swalign() and nwalign(), P_SWNW values between “queryseq” and “databaseseq” are calculated as follows:

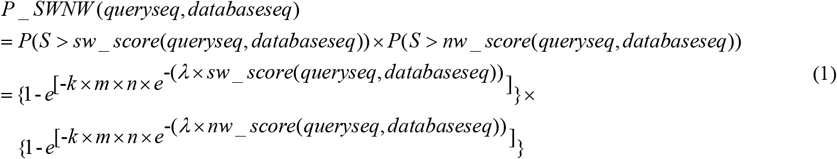

Finally, the SL of “databaseseq” with the minimum out of all P_SWNW is chosen as the predicted SL of “queryseq”.

#### 2.2.4 Definition and Acronyms

FN = False negative number

FP = False positive number

TN = True negative number

TP = True positive number

INN = Input number

ASPS = Amount of sequences passing a preliminary screening by BLAST

ASES = Amount of sequences entering preliminary screening by BLAST

SL= Subcellular localization

SLP = Subcellular localization prediction

SW = Smith-Waterman

NW = Needleman-Wunsch

ACC = Accuracy = (TP+TN)/(TP+FP+FN+TN)

COV = Coverage = ASPS/ASES

SPE = Specificity = TN/(TN+FP)

SEN = Sensitivity = TP/(TP+FN)

## 3. RESULTS

### 3.1 Predictions by BLAST, BLAST+SWNW, and LocTree3 on test Dataset I when the overlap between the test and drill dataset is not removed

When a few sequences in test dataset T1 (1578 out of 1682) also exist in the drill dataset of Dataset I, the prediction accuracy of LocTree3 is 80±3% in SLP for 18 SLs [26]. For such a large overlap, high-version BLAST (from version 2.2.24 to version 2.2.31) can accurately predict SLs for most (1650 out of 1682) test protein sequences with an E-value=1 (see **Table 1**), reaching an average sensitivity of 98.10%. However, low-version BLAST (version 2.2.16) can only reach 94.65% average sensitivity with the same E-value (see **Table 1**). Therefore, a high-version BLAST (version 2.2.31) is used in BLAST+SWNW.

**Table 1.**
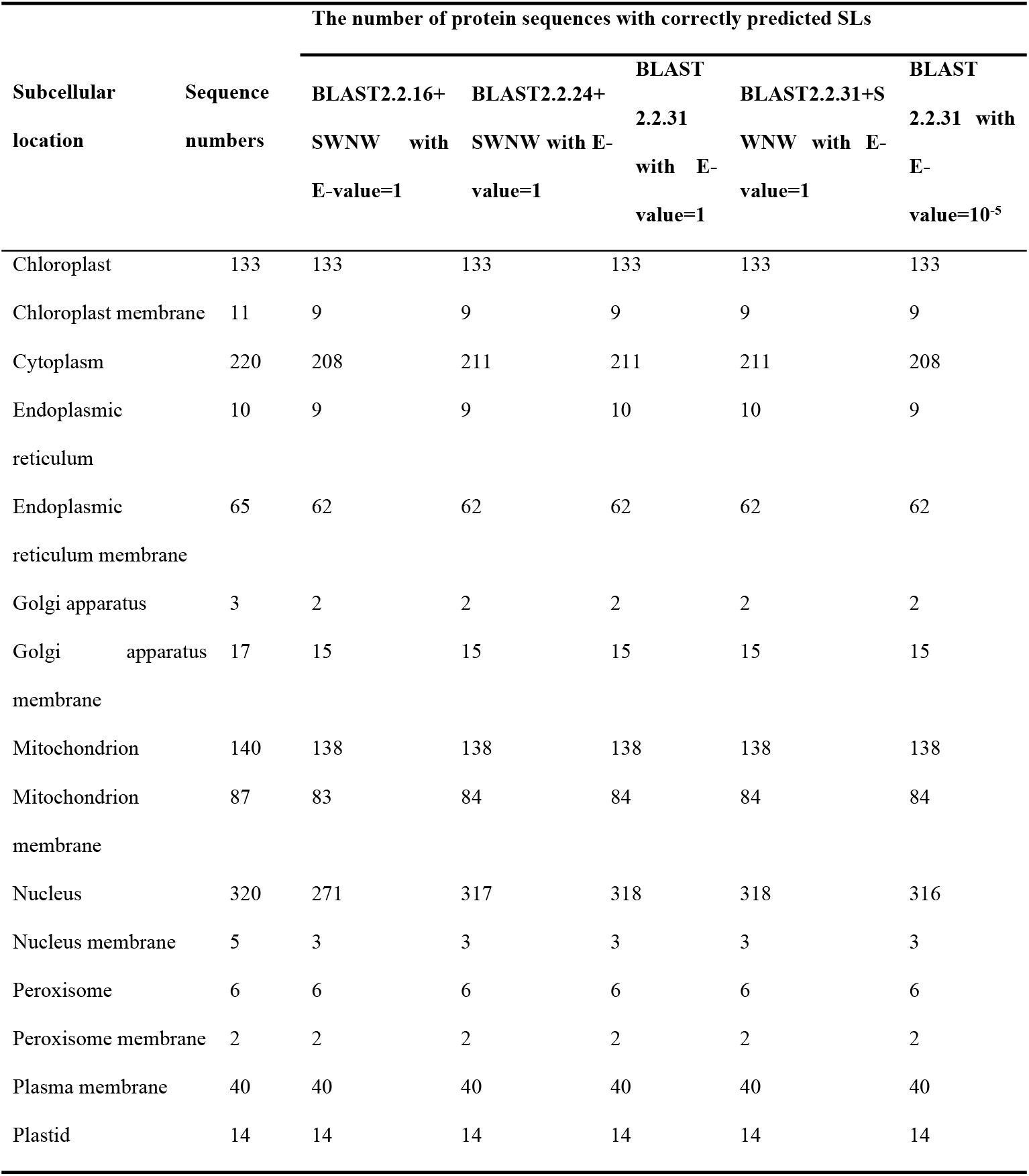
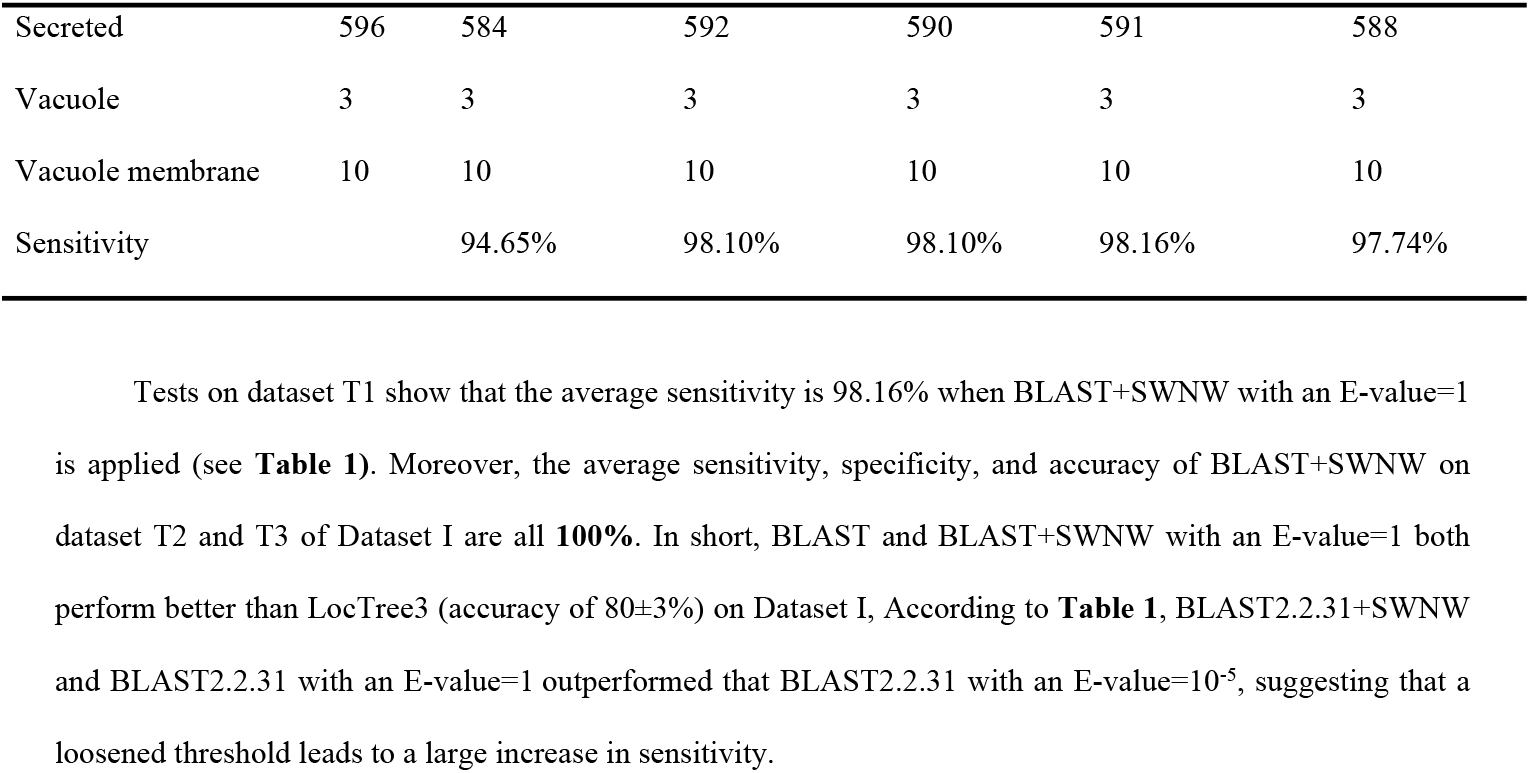
The number of protein sequences with correctly predicted SLs by 5 algorithms on test dataset T1 for 18 SLs when the overlap between T1 and the drill dataset of Dataset I is not removed. (NO need to keep this part)

Tests on dataset T1 show that the average sensitivity is 98.16% when BLAST+SWNW with an E-value=1 is applied (see **Table 1**). Moreover, the average sensitivity, specificity, and accuracy of BLAST+SWNW on dataset T2 and T3 of Dataset I are all **100%**. In short, BLAST and BLAST+SWNW with an E-value=1 both perform better than LocTree3 (accuracy of 80±3%) on Dataset I, According to **Table 1**, BLAST2.2.31+SWNW and BLAST2.2.31 with an E-value=1 outperformed that BLAST2.2.31 with an E-value=10^-5^, suggesting that a loosened threshold leads to a large increase in sensitivity.

### 3.2 Predictions by BLAST and BLAST+SWNW on test Dataset I when the overlap between the test and drill dataset is removed

After the overlapping sequences between test datasets T1, T2, and T3 and the drill dataset of Dataset I were removed, we performed BLAST+SWNW and BLAST on T1, T2 and T3. As described in the Discussion, when we set the E-value to 30, BLAST+SWNW performed best overall in terms of consumed time and sensitivity, so we set 30 as the default E-value if there is no other special instruction. As a result, BLAST+SWNW with an E-value=30 correctly predicts SLs of 2242 sequences, including 1200, 198, and 844 sequences in T1, T2 and T3, respectively, whereas BLAST with an E-value=30 predicts 2218 sequences correctly, including 1167, 201 and 850 sequences in T1, T2, and T3, respectively. Details are shown in S4~7 Table. The average sensitivity, specificity, and accuracy of BLAST+SWNW are 74.51%, 98.52%, and 97.18%, respectively, whereas the average sensitivity, specificity and accuracy of BLAST are 73.71%, 98.47% and 97.09%, respectively; both algorithms outperform LocTree3, which has an accuracy of approximately 80±3% [26].

Additionally, we compared the performance of BLAST+SWNW with pattern recognition algorithms for SLP [26] using KNN, SVM, and random forest on Dataset I with a 168,420-D residue frequency vector representation of protein sequences. However, the results of these algorithms were rather poor (see **S8~10 Table** for details).

### 3.3 Comparison between the predictions of BLAST, BLAST+SWNW, LocTree3, CELLO, and YLOC on Dataset II

In contrast to Dataset I’s 35,738 proteins and 18 SLs, Dataset II consists of 110,168 reviewed mono-localization proteins and 24 SLs and represents a more complete and representative dataset. We further confirmed the advantage of BLAST+SWNW over other algorithms by individually testing BLAST, LocTree3, CELLO, and YLOC on Dataset II.

When we tested BLAST+SWNW with an E-value=30, BLAST+SWNW with an E-value=10^-5^, and BLAST with an E-value=30 on Dataset II using 20-fold cross-validation, the average sensitivity of BLAST+SWNW with an E-value=30 was the highest at **94.47% (104,080/110,168), followed by 94.33% for BLAST** with an E-value=30 and **92.00%** for BLAST+SWNW with an E-value=10^-5^. Realizing a loosened E-value may increase sensitivity, we further tested BLAST+SWNW with E-values varying from 30 to 10^-6^ and confirmed that a higher E-value results in higher sensitivity in this range. Remarkably, the 161 sequences that BLAST with an E-value=30 failed to predict were correctly predicted by BLAST-SWNW, meaning that SW and NW dramatically reinforce BLAST.

Considering the existing differences in SL classifications, we drew a corresponding relationship map of SLs in LocTree3, CELLO, YLOC, and BLAST+SWNW to ensure fairness in their comparisons. For example, the SL “Secreted” in the 18 SLs of Dataset I corresponds to a union of “Secreted” and “Cell wall” in the 24 SLs of Dataset II; SL “Chloroplast” in Dataset I is equivalent to the combination of “chloroplast”, “Thylakoid” and “ Thylakoid membrane” in Dataset II; and SL “Plastid” in Dataset I corresponds to a union of “Plastid” and “Plastid membrane” in Dataset II. In this sense, LocTree3 can predict 22 SLs in Dataset II, which represent all SLs except for “Lysosome” and “Lysosome membrane”. Additional details and the corresponding relationships between the 24 SLs of BLAST+SWNW, 11 SLs of CELLO and 5 SLs of YLOC are shown in **S11 and S12 Table**.

A test dataset with 5497 sequences that were randomly selected from Data II was submitted to the LocTree3 online server, YLOC [14] and CELLO [10]. The input and output file names for LocTree3 are listed in **S13 Table**.

Two out of 24 SLs, “Lysosome” and “Lysosome membrane”, are not available in LocTree3. When we tested sequences from the other 22 SLs, the average sensitivity, specificity and accuracy of LocTree3 were **74.01%, 98.77%, and 97.65%**, respectively, with possible overlaps existing between the drill dataset of LocTree3 and our test dataset. In contrast, even after the overlap was removed, the average sensitivity, specificity, and accuracy of BLAST+SWNW still reached **94.47%, 99.82%**, and 99.60%, and the average sensitivity, specificity, and accuracy of BLAST+SWNW with an E-value=10^-5^ were **92.00%, 99.89%, and 99.57%** respectively based on the same dataset (**S14 Table**). respectively.

We compared the four algorithms (BLAST+SWNW with an E-value=30, BLAST with an E-value=10^-5^, BLAST with an E-value=30 and LocTree3) by applying them to Dataset II, their sensitivities and weighted average sensitivities (marked as “In total”) are shown in **Fig 3**. Obviously, BLAST+SWNW with an E-value=30 and BLAST with an E-value=30 were superior to the other two algorithms. The prediction of 16 of 24 SLs were better with BLAST+SWNW with an E-value=30 than with LocTree3. Undoubtedly, BLAST+SWNW with an E-value=30 will be a better algorithm for predicting protein SLs due to more subcellular locations, higher sensitivity than LocTree3, BLAST, and others.

**Fig 3.**
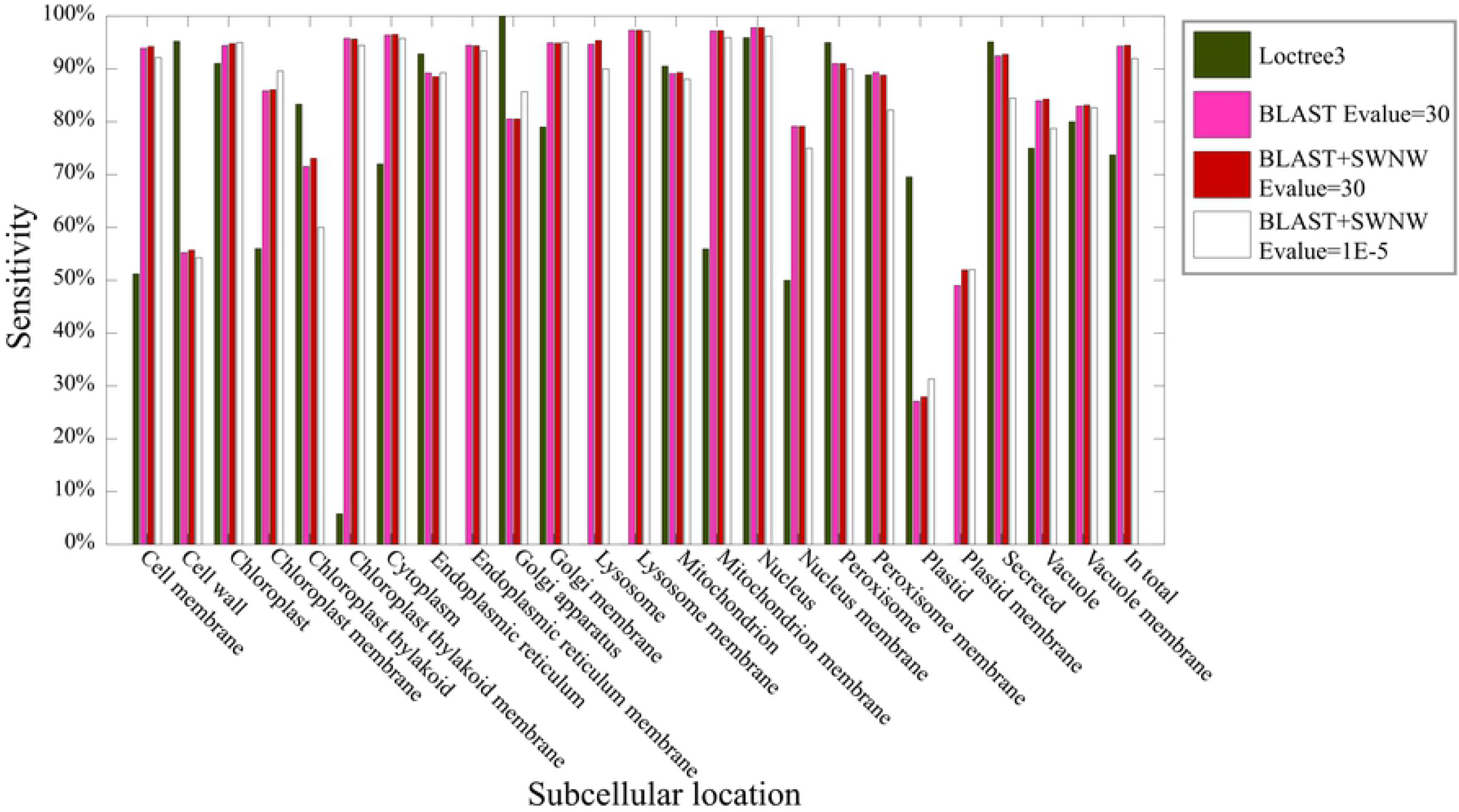
Sensitivities of four algorithms for 24 SLs and their weighted sum sensitivities (marked by “in total”), where the weight of each SL=(ASES of each SL)/(sum of ASES of all SLs).

Based on the sensitivities, specificities, and accuracies listed in **Table 2**, we sort the five algorithms as follows: BLAST+SWNW with an E-value=30 > BLAST with an E-value=30 > BLAST with an E-value=10^-5^ > LocTree3 > CELLO > YLOC.

**Table 2.** Performance comparison between five SLP algorithms on Dataset II. Sensitivities, specificities, and accuracies of CELLO, YLOC, LocTree3, BLAST with an E-value=10^-5^, BLAST with an E-value=30, and BLAST+SWNW with an E-value=30 when tested on Dataset II.

| Method | Sensitivity | Specificity | Accuracy |
| --- | --- | --- | --- |
| CELLO | 58.68% | 96.28% | 93.15% |
| YLOC | 62.96% | 90.74% | 85.19% |
| LocTree3 | 74.01% | 98.77% | 97.65% |
| BLAST with an E-value=10<sup>-5</sup> | 91.88% | 99.89% | 99.57% |
| BLAST with an E-value=30 | 94.33% | 99.81% | 99.59% |
| BLAST+SWNW with an E-value=30 | 94.47% | 99.82% | 99.60% |

### 3.4 Further verification on experimentally annotated rice proteins

SWISS-PROT release 2015_10 contains 3696 reviewed rice proteins. 102 mono-localization rice proteins with published experimental annotations were selected as a new test dataset (S2 Text) from it, Dataset II was used as its drill dataset. we removed the overlap in Dataset II against the 102 sequences, and predicted 102 mono-localization experimental-annotated rice proteins’ SLs by BLAST+SWNW with an E-value=30. As a result, 91 out of 102 proteins were correctly predicted, reaching an average sensitivity 89.22%. (S15 Table).

## 4. DISCUSSION

### 4.1 Optimization of E-values

Next, we determined which E-value increased the sensitivity and reduced the run time of BLAST+SWNW. We randomly selected 1000 proteins (**S3 Text**) from Dataset II as a test dataset, the rest of Dataset II as a drill dataset, and BLAST+SWNW with different E-values to determine the optimal E-value. In **Fig 4**, the time consumed and sensitivity of BLAST+SWNW increased with increasing E-values; however, when the E-value exceeded 30, the time consumed increased sharply while the true positives hardly changed. Thus, we chose 30 as the E-value threshold of BLAST+SWNW.

**Fig 4.**
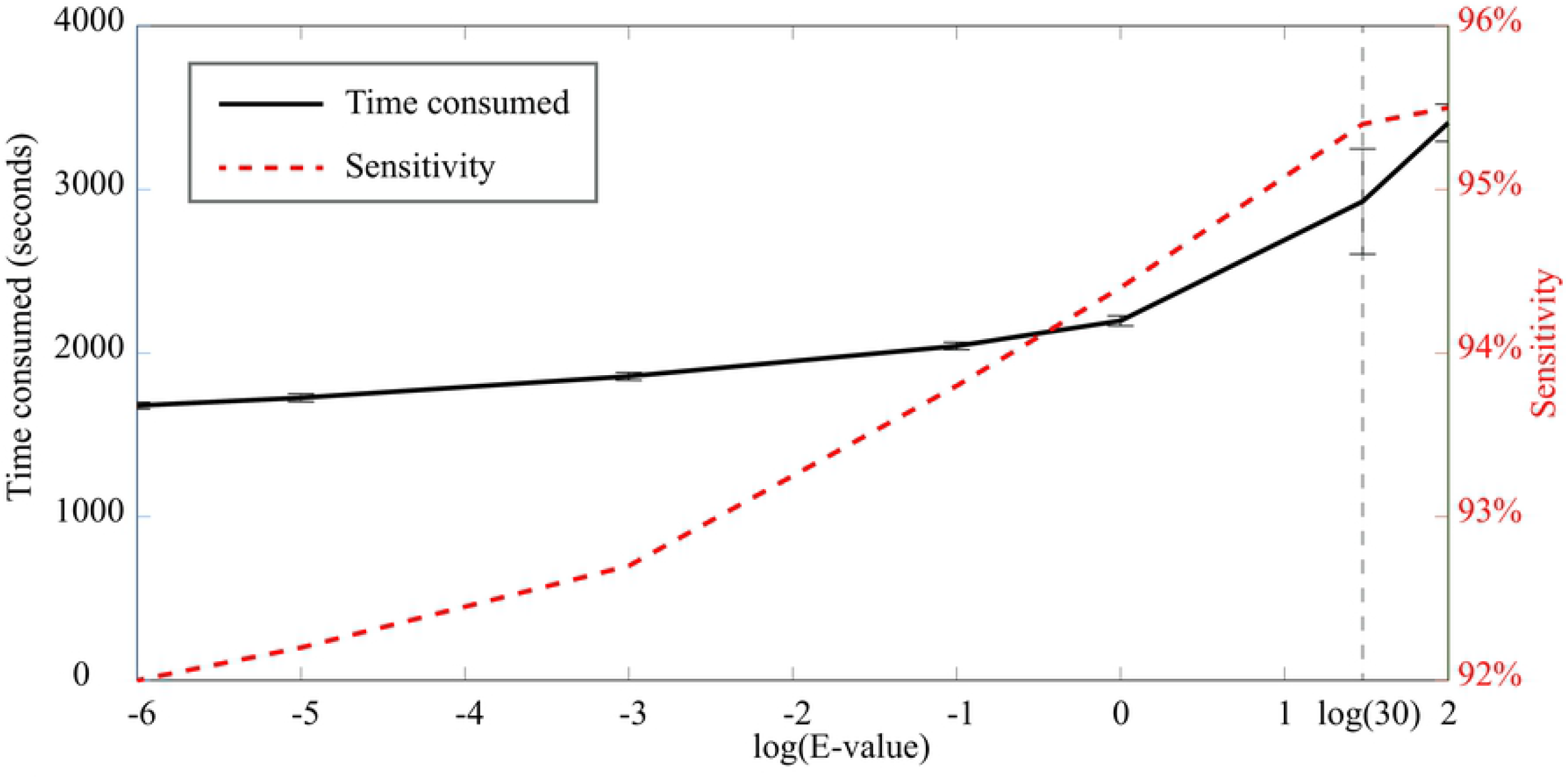
Time consumed and sensitivity of BLAST+SWNW vs. log(E-value) in predicting SLs for 1000 randomly selected proteins.

### 4.2 P_SWNW threshold

Similarly, in order to optimize the threshold of P_SWNW, we selected 10,000 protein prediction results from test Dataset II and drew a line graph representing sensitivity and coverage at different P_SWNW thresholds. The graph shows that the coverage and sensitivity of BLAST+SWNW reach their maximums when the P_SWNW threshold=1 (**Fig 5**). Therefore, 1 was chosen as the default, meaning no threshold was set for SW or NW.

**Fig 5.**
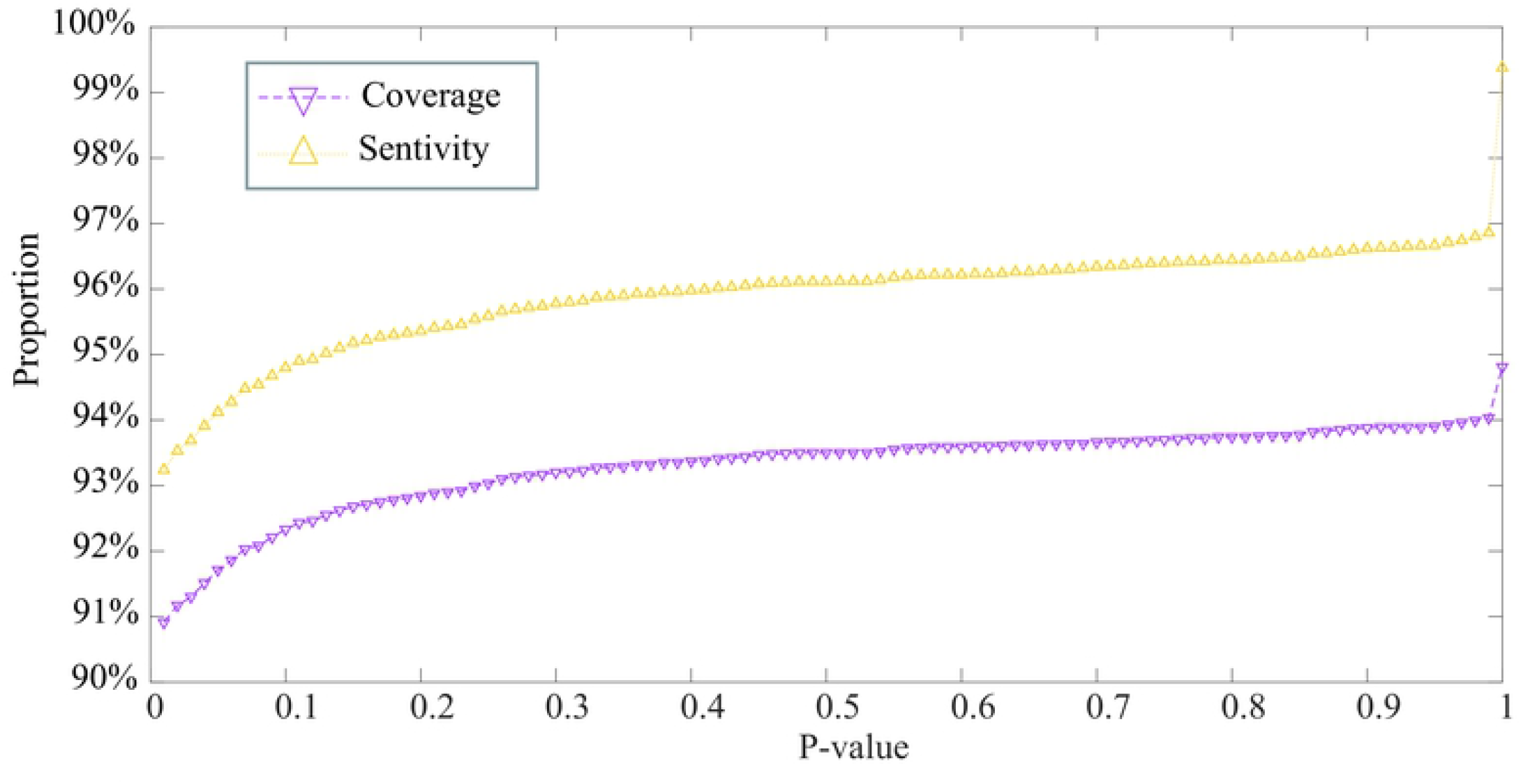
Sensitivity and Coverage of BLAST+SWNW vs. P_SWNW.

### 4.3 The robustness of BLAST+SWNW for different identity between drill and test sequences

After sequences in the drill dataset with 99% or higher identity to test sequences were removed, BLAST+SWNW with an E-value=30 was used on Dataset II and its sensitivity, specificity, and accuracy were 94.56%, 99.81%, and 99.59%, respectively. In addition, when sequences with 95% or higher identity were removed, the sensitivity, specificity, and accuracy were still 93.67%, 99.77%, and 99.51%, 90% and 80% indentity respectively, showing no sign of prediction decline compared to the performance shown in **Table 2**. In summary, even though BLAST, SW, and NW are homology-based algorithms, the excellent results of BLAST+SWNW are not totally dependent on high identity between the test and train dataset.

### 4.4 Reasoning for the effectiveness of BLAST+SWNW

A strategy on hashing and seeking 11-mer for sequences comparison [34] in BLAST is consistent with pattern recognition based on *k*-mer frequencies. Unlike BLAST, the SW and NW algorithms apply dynamic programming to construct a matrix and use backtracking to find an optimal alignment between two given sequences. In this sense, BLAST can be more strongly reinforced by SW and NW than by pattern recognition algorithms. In other words, BLAST+SWNW is more powerful than BLAST+pattern recognition, such as LocTree3. Additionally, the poor performance of pattern recognition algorithms (shown in **S1 Text, S8~10 Table**) suggests that additional intrinsic features should be added and used in SLP to improve prediction sensitivity, specificity, and accuracy.

## 5. ACKNOWLEDGEMENT

We thank Dr. Hong Yu for his valuable comments to the earlier version of this manuscript. We are grateful to Prof. Yidan Ouyang for her precious suggestions and long-term help.

